# Protein restriction amplifies nucleus accumbens dopamine responses to protein-containing food during operant feeding in male mice

**DOI:** 10.64898/2026.07.31.741998

**Authors:** Hamid Taghipourbibalan, Stefan W Huijgens, K. Linnea Volcko, James E. McCutcheon

## Abstract

Animals defend protein intake strongly, yet how protein status shapes moment-to-moment reward signalling in mesolimbic circuits remains unclear. Here, we examined how dietary protein restriction shapes nucleus accumbens (NAc) core dopamine signalling during operant feeding in the home-cage. C57BL/6 mice expressing the GRAB_DA_ sensor in NAc core performed 1 h fixed-ratio 1 (FR1) and progressive ratio (ProgRatio) sessions to earn grain or sucrose pellets across three dietary phases; an initial non-restricted phase (NR1), a protein-restricted phase (PR), and a return to the non-restricted diet (NR2). Home-cage food intake remained stable across phases, whereas bodyweight gain was markedly reduced during PR and partially recovered in NR2. Behaviourally, operant responding increased during PR under both FR1 and ProgRatio schedules, with the most robust enhancement observed for grain pellets. Fibre photometry revealed a more selective neural effect. During FR1, pellet delivery evoked a larger NAc dopamine response for grain than for sucrose specifically during PR, whereas no pellet-type difference was detected during NR1 or NR2. By contrast, under progressive ratio, dopamine responses at pellet delivery and pellet retrieval did not differ between pellet types in any dietary phase. Together, these findings show that protein restriction does not globally amplify reward-related behaviour or dopamine signalling, but instead selectively biases NAc dopamine encoding in a manner that depends on nutritional status of the animal, nutrient context, and task demands.

## 1. Introduction

Animals tightly regulate their intake of protein, often at the expense of overall energy balance. This is formalised as the protein leverage hypothesis (PLH), which suggests protein intake is defended more strongly than other macronutrients, such that dilution of dietary protein drives compensatory overeating to restore a protein target (Raubenheimer & Simpson, 2023; Simpson & Raubenheimer, 2005). In rodents, geometric analyses of macronutrient choice show that limiting protein shifts intake towards protein-rich options and alters adiposity and metabolic health (Hill et al., 2019a; Morrison & Laeger, 2015; Sørensen et al., 2008). These findings position protein appetite as a key organising principle in the control of food intake and raise the question of how internal protein status reshapes the activity of neural circuits that govern reward, motivation, and choice.

In rodent models, behaviour towards macronutrients is changed markedly by protein restriction. Protein-restricted mice and rats develop strong preferences for protein-containing foods and show elevated palatability for such options (Hill et al., 2022; Murphy et al., 2018; Naneix, Peters, & McCutcheon, 2020; Volcko & McCutcheon, 2022; Wu et al., 2024), suggesting that internal protein need is translated into altered reward valuation. Concordant with these findings, protein restriction in mice and rats increases motivation for protein containing food in operant paradigms (Chiacchierini et al., 2022; Khan et al., 2025).

Mesolimbic dopamine signalling is thought critical for shaping food valuation, effort expenditure, and macronutrient choice (Betley et al., 2013; Keen-Rhinehart et al., 2013) and so may underpin the behavioural changes after protein restriction. In rats, protein restriction alters dopamine release in the NAc in an age-dependent manner (Naneix, Peters, Young, et al., 2020) and biases ventral tegmental area (VTA) activity during choices between protein-rich and non-protein options (Chiacchierini et al., 2021). In mice, similar effects are found with protein restriction biasing dopamine responses towards intraoral protein rather than carbohydrate (Khan et al., 2025). In addition, protein restriction reduces dopamine responses to sucrose (Wu et al., 2024). These effects on dopamine signalling have been linked to endocrine changes induced by protein restriction, notably an increase in fibroblast growth factor 21 (FGF21) (Hill et al., 2019b, 2022; Laeger et al., 2014). As such, several behavioural and neural changes associated with protein restriction are abolished if FGF21 is absent either globally or centrally (Hill et al., 2019a, 2020; Khan et al., 2025).

To date, the effects of protein restriction on dopamine circuits have not been assayed during ongoing operant behaviour that discriminates motivation for different macronutrients. To address this, we tested how dietary protein restriction alters dopamine dynamics during operant feeding with different effort requirements. We used fibre photometry to measure dopamine release in NAc core of mice performing fixed and progressive ratio tasks for pellets that differed in protein content. NAc core was targeted as this region is implicated in translating reward-related information into goal-directed and effortful actions (Salamone & Correa, 2024). We show that protein restriction leaves non-effortful intake largely unchanged while altering operant behaviour and reorganising dopamine signalling in a task- and nutrient-dependent manner.

## 2. Materials and Methods

### 2.1 Animals and housing

Twelve adult male C57BL/6NRj mice (6–8 weeks old, 21–30 g; Janvier Labs, France) were used. Animals were acclimatised for 5 days in a temperature-(22 ± 0.5 °C) and humidity-(56 ± 2%) controlled facility maintained on a 12 h light/dark cycle (lights on at 12:00 AM; lights off at 12:00 PM). After acclimation, mice were moved to the colony room with identical environmental conditions. Mice were housed two per cage in Eurostandard type II L cages (1284 L; Techniplast; 365 × 207 × 140 mm; 530 cm^2^ floor area) modified to accommodate Feeding Experimentation Devices (FED3). A FED3 unit is a programable and automated home-cage feeding device that dispenses 20 mg food pellets and accurately logs feeding events such as pellet delivery, retrieval and nose pokes in different modes of activity (Matikainen-Ankney et al., 2021).The mice were separated by a perforated divider that permitted visual, olfactory, and limited tactile communication while preventing major physical contact. These cages had two lateral ports designed for mounting FED3 devices; outside photometry recording rounds, the ports were covered with removable 3D printed panels (Fig. S1A) and during recording sessions, the FED3 devices were mounted (Fig. S1B-C). The cages contained approximately 1 kg of wood-shaving bedding, and each mouse received 3.65 g of cotton nesting material. Water and standard chow (Ssniff Rat/Mouse Maintenance Diet) were available ad libitum unless otherwise stated. All procedures complied with EU Directive 2010/63/EU and were approved by the Norwegian Food Safety Authority (FOTS #30889).

### 2.2 Surgical procedures

Surgical procedures followed those previously described (Volcko et al., 2025), with adjustments for GRABDA expression in the NAc core. Mice were anesthetised with 1.5-2 % isoflurane in air and secured in a stereotaxic frame (Kopf Instruments, USA). Pre-operative systemic analgesia was provided with buprenorphine (0.1 mg/kg, s.c.) and meloxicam (5 mg/kg, s.c.), along with local anaesthesia using bupivacaine (1 mg/kg, s.c.). A craniotomy was made above the NAc core (coordinates relative to Bregma, flat skull: AP +1.5, ML +1.0, DV −4.65). A total of 500 nL AAV1-hSyn-GRAB_^DA^_2m was infused at 100 nL/min through a 33 G Nanofil syringe coupled to a UMP3 microsyringe pump (World Precision Instruments, USA). The needle remained in place for 10 min to allow viral diffusion. A fibre-optic cannula (1.25 mm ferrule, 400 ^µ^m core, 0.48 NA; RWD) was then implanted just dorsal to the injection site (DV −4.55) and secured with dental cement (Super-Bond C&B) (Fig1. C). Following surgery, animals received meloxicam for two additional days and were monitored daily for one week to ensure full recovery. Behavioural testing commenced no earlier than 21 days after surgery, allowing sufficient time for postoperative recovery and GRABDA sensor expression.

### 2.3 Experiment paradigm and behavioural setup

The experiment comprised of three consecutive dietary phases: an initial non-restricted phase (NR1; 20% casein: D11051801; Research Diets), a protein-restricted phase (PR; 5% casein: D15100602; Research Diets), and a return to the non-restricted phase (NR2; 20% casein). Each phase lasted two weeks. During the first week of each phase, mice remained in their home cages with ad libitum access to the assigned maintenance diet. Behavioural and neural recordings were performed during the second week of each phase while animals remained on the same diet. Animals were switched to the next dietary phase after completion of the recording week (Fig. 1A).

**Figure 1.**
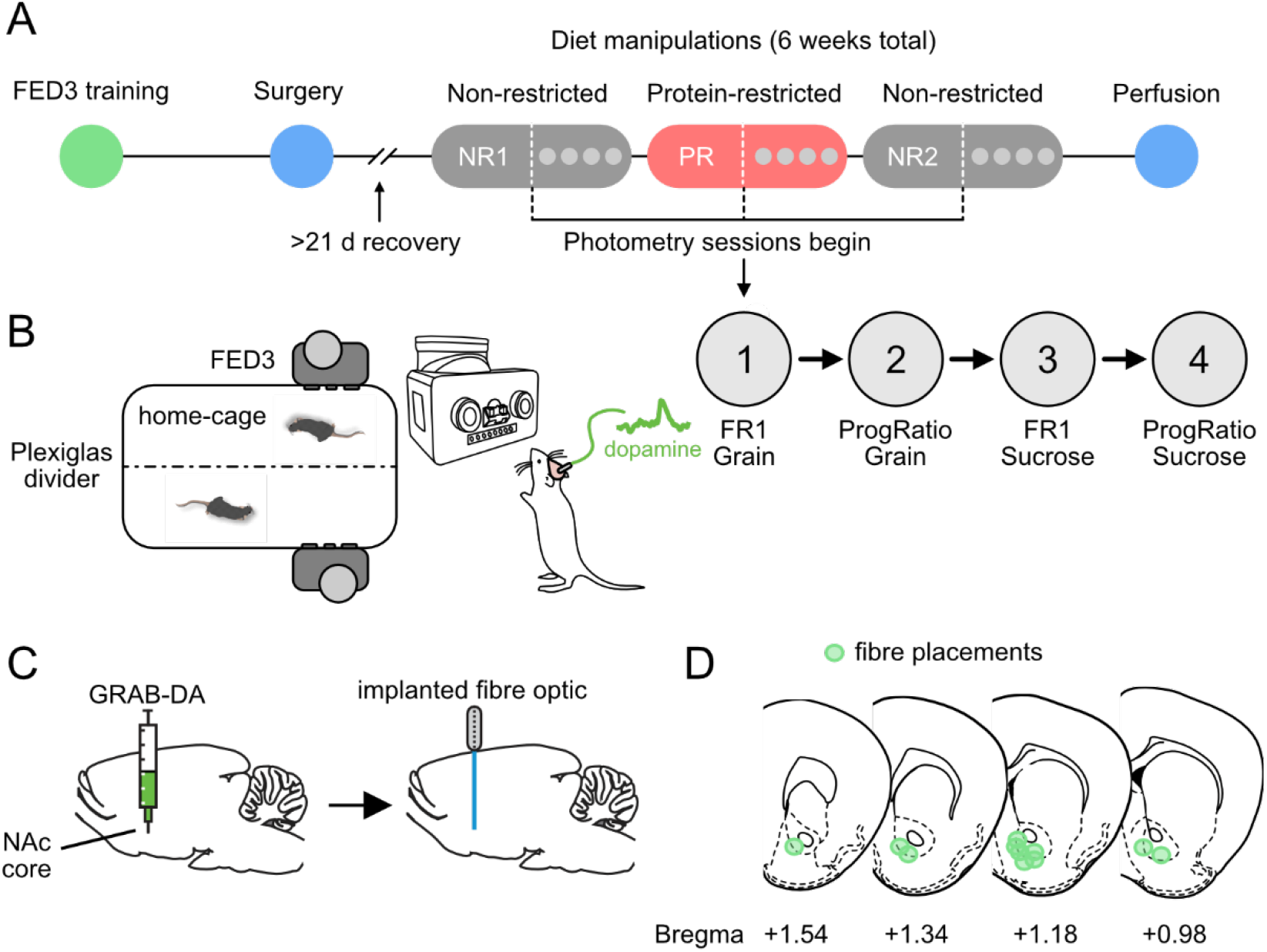
Schematic showing experimental timeline and methods. (A) Timeline showing progression of mice through training, surgery, and dietary manipulations. In each diet phase, operant sessions began after 1 week on diet and two sessions with grain pellets (FR1 and progressive ratio) and two sessions with sucrose pellets (FR1 and progressive ratio). At the end of experiment, mice were perfused for confirmation of fibre placement and virus expression. (B) Mice were contact-housed, two per cage, with a Plexiglas divider. FED3 devices were attached to the side of the cage for training and performance of operant behaviours. When not in use a removable plastic cover was used. (C) Mice were prepared for fibre photometry recordings by injecting GRAB-DA into nucleus accumbens (NAc) core and implanting a fibre optic. (D) Fibre placements were located in NAc core.

#### 2.3.1 Home-cage behavioural and recording setup

Before the experimental recording rounds, mice were trained in their home cages to collect 20 mg grain pellets (#F0163; Bio-Serv) from FED3 devices, first in free-feeding mode for 2 days and then under fixed-ratio 1 (FR1) and closed-economy modes for 1 day each. In free-feeding mode, FED3 dispenses a food pellet immediately after the previous pellet is retrieved, ensuring that food remains continuously available without any operant response requirement. In the FR1 schedule, a single left nose poke (active poke) results in delivery of one food pellet. In the closed-economy mode, the response requirement increases progressively with each pellet earned, such that an increasing number of nose pokes is required to obtain successive pellets. However, unlike a standard progressive-ratio schedule, the response requirement resets to one poke if the animal remains inactive for 30 min. This schedule is used during training to allow extended access to pellets without mice reaching excessively high response requirements that would otherwise limit pellet acquisition over the course of a full day. Moreover, to reduce novelty during subsequent testing, mice were also given 20 mg grain and sucrose pellets (#F07595; Bio-Serv) in a dish in the home cage. In addition, animals were habituated to transport to the recording room and to connection of the fibre optic patch cable.

For recording sessions, the cage food hopper was emptied 3 h before each session and mice were moved in their home cages from the colony room to the recording room 10 min before session onset. At the start of each recording week, the removable side panels of the home cages were replaced with FED3 devices, which remained mounted throughout the week but were switched on only during recording sessions. Mice were then connected to the fibre optic cable and allowed to freely explore the cage and interact with their individual FED3 devices (Supplementary Video 1). Behavioural testing was performed in the animals’ customized home cages using FED3 devices (Fig. 1B) and the RTFED (PiTTL) system for real-time behavioural monitoring, as described in detail in (Taghipourbibalan & McCutcheon, 2026b).

The RTFED system logged behavioural events, including left and right nose-pokes, pellet onset (delivery to the pellet well), and pellet offset (retrieval by the animal), and relayed these events to an RZ10x system running Synapse software (Tucker-Davis Technologies, Alachua, FL, USA). The TDT system controlled two LED light sources, one blue (465 nm) and one violet (405 nm). The blue light was sinusoidally modulated at 330 Hz and the violet light at 210 Hz. A fluorescence minicube (Doric Lenses, Quebec, Canada) combined the two excitation wavelengths. Light was delivered via a 400 μm optical patch cord (Doric Lenses) connected to the implanted ferrule using a black ceramic sleeve (RWD), and the same optical path was used to collect emitted fluorescence from the brain.

#### 2.3.2 Fibre photometry recording sessions

During the recording week of each dietary phase, mice underwent four 1h fibre photometry sessions conducted within the first 3 h of the dark phase in a recording room under dim red illumination. These sessions consisted of FR1 grain, Progressive Ratio (ProgRatio) grain, FR1 sucrose, and ProgRatio sucrose, using 20 mg pellets (Fig. 1B, Right panel). Additional sessions using 35% casein pellets were also performed during each recording round, but these were exploratory pilot sessions and are not included in the present report. The macronutrient composition of the maintenance diets and test pellets is provided in Table S1.

### 2.4 Histological procedure

At the end of the experiment, brains were collected to verify viral expression and fibre placement in the NAc core. Only animals with correct placements and sufficient viral expression were included in the final analysis (Fig. 1D). Mice were deeply anaesthetised with 0.3 mL ZRF mix (zolazepam 3.3 mg/mL; tiletamine 3.3 mg/mL; xylazine 0.45 mg/mL; fentanyl 2.6 μg/mL) and transcardially perfused with heparinised saline followed by 4% paraformaldehyde (PFA). Brains were extracted, post-fixed overnight in 4% PFA, and transferred to 30% sucrose containing ProClin 150 (Sigma, 49376-U) for cryoprotection. Coronal sections (40 μm) were cut using a Leica microtome (Leica, Deer Park, IL, USA). Every third section was mounted onto Superfrost Plus slides and coverslipped. Native fluorescence of the GRAB_^DA^_ virus was used to confirm the location and spread of the injection (Fig. 1D), and the fibre tract was inspected to assess implant position relative to the NAc core (target coordinates: AP +1.5, ML +1.0, DV -4.55 to -4.65).

### 2.5 Data processing and statistical approach

Behavioural and fibre photometry data were processed using custom scripts written in Python and R (v4.4.1) on a 64-bit Windows platform. Across analyses, repeated measurements from the same animals were treated using within-subject statistical designs. Descriptive statistics are reported as mean ± SEM unless otherwise stated. For post hoc testing, p values were adjusted using the Holm method within each family of comparisons.

#### 2.5.1 Behavioural data

Home-cage food intake and body weight were recorded manually across the three dietary phases. For each mouse, food consumed during each phase was calculated from hopper measurements, and bodyweight change was calculated across the same phase. Bodyweight change was expressed as percentage change for the main home-cage phase analysis. The effects of dietary phase on food intake and bodyweight change were assessed using one-factor repeated-measures ANOVA with phase as the within-subject factor. Where appropriate, paired post hoc comparisons between phases (NR1 vs PR, PR vs NR2, and NR1 vs NR2) were performed with Holm correction.

Operant behavioural measures were extracted from raw FED3 files using custom scripts in Python. For each session, the main parameters including pellet count, left (active) poke count, and, for progressive-ratio sessions, the maximum completed ratio (i.e., breakpoint) were extracted from the raw FED3 data. Analyses were performed on sessions with either grain or sucrose pellets performed under FR1 and ProgRatio schedules during each diet phase (NR1, PR, and NR2. For operant behavioural outcomes, repeated-measures ANOVA with diet phase and pellet type as within subject factors were conducted. Holm-corrected post hoc tests were conducted if the interaction term was significant. Mice that did not earn any pellets or perform any pokes were excluded from analysis. This resulted in exclusion of 2 mice from FR1 analysis and 3 mice from ProgRatio analysis.

As a supplementary analysis, we examined whether inter-individual variation in FR1 responding during the 1 h operant sessions was associated with home-cage food intake or bodyweight change measured during the corresponding dietary phase. For this purpose, mean pellet counts from the FR1 phase were matched by mouse and phase to home-cage food consumption and bodyweight change, and Pearson correlation coefficients were calculated separately for grain and sucrose within each phase.

#### 2.5.2 Fibre photometry data

Raw fibre photometry recordings were acquired as dual-wavelength signals and analysed in Python using custom scripts. For each session, the activity-dependent blue channel was processed together with the simultaneously recorded UV isosbestic channel to generate a motion- and bleaching-corrected signal (Konanur et al., 2020). Behavioural events were identified from TTL timestamps recorded alongside the photometry data, and peri-event signal segments were extracted from 5 s before to 10 s after each event. Analyses focused on the time of pellet delivery, which corresponds to with active pokes in FR1 sessions, and with the final poke completing a ratio in ProgRatio sessions. We report analysis of pellet retrieval in Supplemental Material, but as latency between pellet delivery and retrieval was typically very short, these analyses look very similar.

Because the number of available events varied across mice, phases, pellet types, and schedules, photometry analyses were performed at the mouse level to avoid pseudoreplication. Within each mouse and condition, only the first *K* events were retained before averaging. The value of *K* was chosen empirically on the basis of an exploratory event-availability analysis designed to maximise inclusion of mice across conditions while reducing imbalance in event counts contributing to the signal averages. For the final analyses, *K* was set to 6 events per mouse for FR1 left-poke and FR1 pellet-offset analyses, and 4 events per mouse for ProgRatio pellet-onset and pellet-offset analyses.

For each mouse, the retained event-aligned traces were averaged and then baseline-corrected by subtracting the mean signal in the -2 to 0 s pre-event window. The response metric was the area under the curve (AUC) during the 0 to 5 s post-event window. Statistical analyses were performed on these mouse-level values. One-way repeated-measures ANOVA was used to test whether responses to grain and sucrose pellets differed. For data visualization, average traces were plotted with bootstrap confidence intervals, alongside paired mouse-level summary plots of the corresponding 0-5 s AUC values. The difference between the AUC of grain pellets and sucrose pellets was calculated and this was compared against zero (no difference between pellets) using a Bonferroni-adjusted one-sample t-test and between phases using unpaired t-test. A more detailed description of photometry preprocessing, event selection, and exclusion handling is provided in the Supplementary Materials (section S2.5.2).

### 2.9 Code and data availability

All analysis code is available at https://github.com/Htbibalan/photofed_paper.git and raw and processed data files are available at https://doi.org/10.5281/zenodo.21024650.

## 3. Results

### 3.1 Protein restriction alters body-weight gain without affecting non-FED intake of large food pellets

Total food intake of large chow pellets from the standard cage hopper did not differ significantly across dietary phases (Fig. 2A). Mean intake was similar during NR1, PR, and NR2. A repeated-measures ANOVA revealed no significant effect of phase on food intake (F_2,22_ = 0.92, p = .413), and all paired post hoc comparisons were non-significant (Holm-corrected p ≥ .523). In contrast, body-weight change differed significantly across phases (Fig. 2B). Animals gained weight during NR1 (one-sample t-test vs. zero, p < .001), did not show weight gain during PR (p = .191), and resumed weight gain during NR2 (p < .001). Repeated-measures ANOVA confirmed a significant effect of phase on body-weight change (F_2,22_ = 13.47, p < .001). All pairwise phase comparisons were significant after Holm correction (NR1 vs PR: p = .002; PR vs NR2: p = .018; NR1 vs NR2: p = .048).

**Figure 2.**
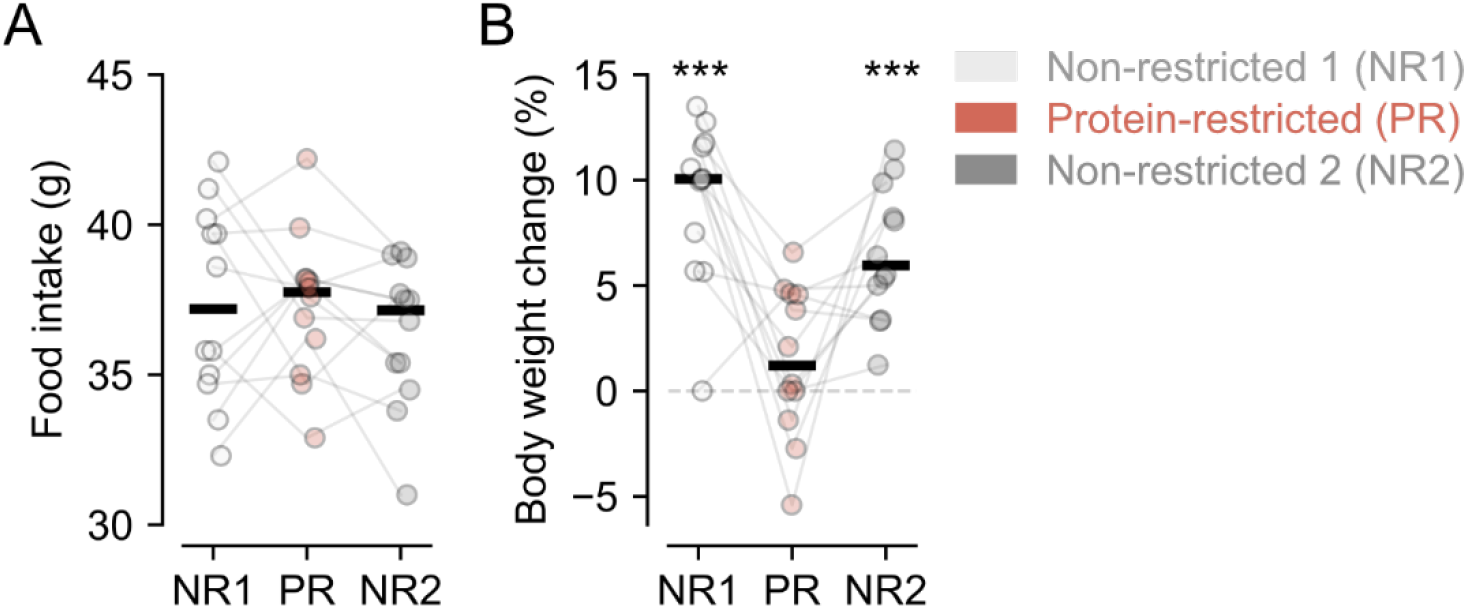
Mice did not change food intake when protein-restricted but body weight gain was reduced. (A) Food intake was the same in mice across all diet phases. (B) Body weight increased while on non-restricted diet but did not increase on protein-restricted diet. Thick black lines are mean and connected circles are individual mice. ***, *p* < 0.001 and ns, non significant vs. zero.

Dietary protein restriction therefore did not induce compensatory hyperphagia at the level of gross food intake. Despite similar total food intake across phases, protein restriction markedly suppressed bodyweight gain. To note, these measurements reflect the changes occurring across each phase (two weeks) as a whole; more fine-grained within-phase analyses may reveal temporal dynamics not captured by the phase-level summary(Taghipourbibalan & McCutcheon, 2026a).

### 3.2 Dietary phase differentially modulates operant responding across reinforcement schedules

Under FR1 conditions, total pellets earned varied significantly across dietary phase for both grain and sucrose pellets (Fig. 3A). As such, there was a main effect of Phase (F_2,18_ = 24.92, p < .001), a main effect of Pellet Type (F_1,9_ = 6.93, p = .027), and a significant interaction (F_2,18_ = 5.01, p = .019). Holm-adjusted post hoc tests showed that across diet phases, mice took more grain pellets in PR than in either NR phase (PR vs NR1: p = .008; PR vs. NR2: p < .001) while there was no difference in number of grain pellets between NR1 and NR2 (p = .406). Mice took more sucrose pellets in PR than in NR2 and there was a trend towards more sucrose pellets in PR than in NR1 (p = .081). Mice also took more sucrose pellets in NR1 than NR2 (p = .002). When comparing within each phase, post hoc tests revealed that mice took more grain pellets than sucrose pellets in PR (p = 0.18) and in NR2 (p = .007), but a similar number of each in NR1 (p = .680).

**Figure 3.**
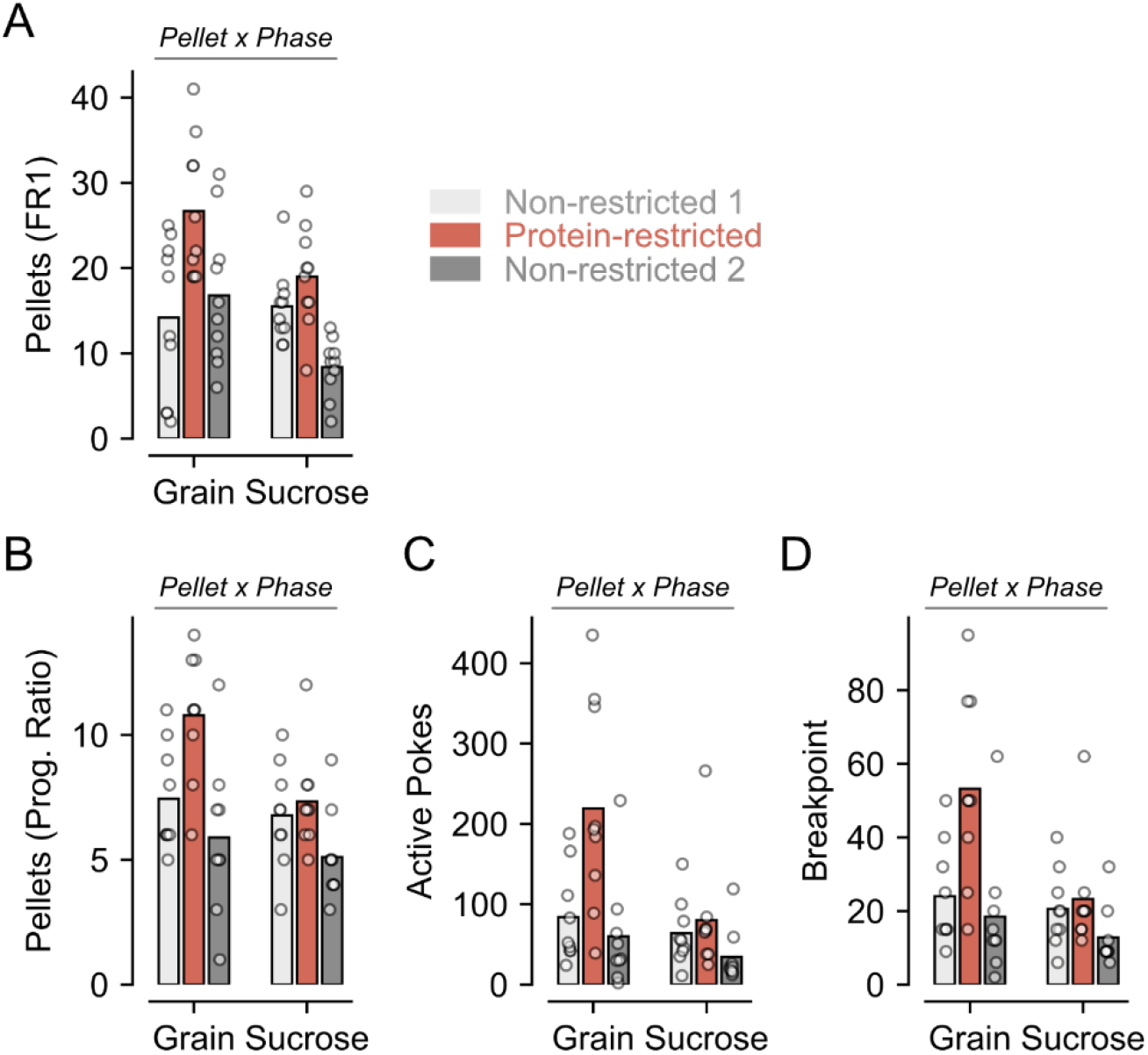
Mice took more pellets and worked harder for food when protein-restricted than non-restricted. (A) On a fixed ratio 1 (FR1) schedule where each active poke results in delivery of a pellet, mice earned more pellets when protein restricted than non-restricted. This effect was driven by differences in grain pellets rather than sucrose pellets. (B) On a progressive ratio schedule where each subsequent pellet requires completion of more nose pokes than the last one, mice earned more grain pellets when protein-restricted than non-restricted. (C) Mice made more active pokes for grain pellets when protein restricted than non-restricted and more active pokes for grain pellets than sucrose pellets when protein restricted. (D) Mice reached higher breakpoints when responding for grain pellets than sucrose pellets when protein restricted. Significant interactions between pellet type (grain vs. sucrose) and diet phase (NR1 vs. PR vs. NR2) were found for all measures.

Under ProgRatio, number of pellets earned also varied significantly across phase for both pellet types (Fig. 3B). Specifically, there was a main effect of Phase (F_2,16_ = 11.77, p < .001), a trend towards an effect of Pellet Type (F_1,8_ = 4.29, p = .072), and a significant interaction (F_2,16_ = 3.70, p = .048). Holm-adjusted post hoc tests showed that across diet phases, mice earned more grain pellets in the PR phase than in either NR phase (PR vs NR1: p = .027; PR vs. NR2: p = 0.027). There was trend towards mice earning more grain pellets in NR1 than in NR2 (p = .071). Mice earned more sucrose pellets in PR phase than in NR2 phase (p = .002) but earned a similar number when other phases were compared (PR vs NR1: p = .567; NR1 vs NR2: p = .141). When comparing within each phase, during the PR phase mice earned more grain pellets than sucrose (p = .008) but earned similar numbers of grain and sucrose pellets within each NR phase (NR1: p = .428; NR2: p = .553).

Active pokes were also analysed and varied significantly across phase for both pellet types (Fig. 3C). There was a main effect of Phase (F_2,16_ = 10.13, p = .008), a main effect of Pellet Type (F_1,8_ = 5.79, p = .043), and a significant interaction (F_2,16_ = 4.91, p = .042). Holm-adjusted post hoc tests showed that across diet phases, mice performed more active pokes for grain pellets in the PR phase than in either NR phase (PR vs NR1: p = .045; PR vs. NR2: p = .045). There was no difference in active pokes for grain between NR1 and NR2 (p = .159). For sucrose pellets, mice performed more pokes in PR phase than in NR2 phase (p = .035) but made the same number of pokes when other phases were compared (PR vs NR1: p = .571; NR1 vs NR2: p = .196). When comparing within each phase, during the PR phase mice made more pokes for grain pellets than for sucrose (p = .026), but pokes did not differ between pellets in either NR phase (NR1: p = .374; NR2: p = .376).

Breakpoints (maximum pellet cost) reached under ProgRatio showed a similar overall pattern with differences across phase and between pellets (Fig. 3D). As such, there was a main effect of Phase (F_2,16_ = 10.94, p = .001), a main effect of Pellet Type (F_1,8_ = 5.57, p = .046), and a significant interaction (F_2,16_ = 5.45, p = .016). Mice reached higher breakpoints in PR than in NR and breakpoints for grain pellets were higher than for sucrose pellets. Holm-adjusted post hoc tests showed that across diet phases, mice reached higher breakpoints for grain pellets in the PR phase than in either NR phase (PR vs NR1: p = .033; PR vs. NR2: p = .033). There was no difference in breakpoints for grain between NR1 and NR2 (p = .245). For sucrose pellets, mice reached a higher breakpoint in PR phase than in NR2 phase (p = .014) but had similar breakpoints between other phases (PR vs NR1: p = .660; NR1 vs NR2: p = .121). When comparing within each phase, during the PR phase the breakpoint for grain was greater than for sucrose (p = .017) but did not differ between pellets in either NR phase (NR1: p = .524; NR2: p = .419).

Together, these findings show that protein restriction increases operant responding in both a low effort scenario (FR1) and when demands are increased (progressive ratio), with a more robust enhancement when mice were working for grain pellets (which contain protein) than for sucrose pellets (which do not). In addition, differences in responding for the pellet types was not observed in the first non-restricted phase (NR1) but were observed in the second non-restricted phase (NR2), suggesting that the effects of protein restriction on pellet preference and associated motivation persist even after restoration of protein levels.

### 3.3 Inter-individual variability in operant responding is not explained by energy balance

To assess whether inter-individual differences in operant responding were related to broader energy-balance measures, we examined the relationship between behavioural performance during recording sessions and bodyweight/food intake measures collected in the home cage during the same dietary phase. These analyses focused on FR1 performance, where pellet count provides a direct index of session intake, and tested whether mean pellets earned was associated with food consumption across each phase or bodyweight change across animals. Pearson correlation analyses were performed separately for each dietary phase and pellet type. No significant correlations were detected between FR1 pellet count and either home-cage food intake or bodyweight change in any phase for either grain or sucrose pellets (all p > 0.05; Fig. S2). Correlation coefficients were generally weak to moderate in magnitude and varied in direction across conditions, indicating no consistent relationship between passive intake measures and operant responding. For example, the largest trend was observed for Sucrose pellets while protein-restricted, where FR1 pellet count showed a moderate negative correlation with phase food intake, but this effect did not reach significance (r = -.529, p = .0767). Overall, these findings indicate that inter-individual variability in FR1 responding was not explained by differences in overall food intake or bodyweight change across the corresponding dietary phase. These results suggest that the behavioural effects of dietary protein manipulation during recording sessions with FED3 devices were dissociated from gross measures of energy balance. Thus, the state- and phase-dependent changes in operant responding observed across dietary transitions are unlikely to be explained simply by how much food animals consumed in the home cage or by concurrent changes in bodyweight and instead are more consistent with altered motivational valuation of the available pellets.

### 3.4 FR1 (low-effort intake): protein restriction selectively enhances pellet-specific NAc dopamine responses

Under FR1 conditions, NAc dopamine responses discriminated between pellet types specifically during the protein-restricted phase (Fig. 4A-B). For pellet delivery–aligned signals, the 0-5 s post-event AUC (Fig. 4C) showed a significant effect of pellet type in PR, with grain evoking a larger dopamine response than sucrose (F_1,11_ = 21.67, p < .001; Holm-corrected post hoc p = .002). In contrast, no significant pellet-type difference was detected in NR1 (F_1,8_ = 0.06, p = .811) or NR2 (F_1,7_ = 1.00, p = .350). These results were confirmed by calculating the difference between the dopamine AUC associated with each type of pellet and comparing it to zero (equal AUC between pellets; Fig. 4D). This analysis showed that grain pellets were associated with greater dopamine during PR (Bonferroni-corrected one-sample t-test; t_11_ = 4.96, p = .001) but not during NR1 (t_7_ = 0.24, p = 1.0) or NR2 (t_6_ = 0.75, p = 1.0). A similar pattern was observed for pellet retrieval–aligned signals (Fig. S3). In addition, we compared latencies from pellet delivery to pellet retrieval (Fig. S4) and found that these were typically very short (94.54% FR1 latencies < 5 s). During the NR1 phase, latencies to retrieve sucrose pellets were shorter than those to retrieve grain pellets (unpaired t-test: t_17_ = 2.13, p = .048), but latencies did not differ between grain and sucrose in either the PR phase (t_22_ = 0.12, p = .903) or the NR2 phase (t_16_ = 1.22, p = .240). Together, these data show that under low-effort FR1 conditions, protein restriction selectively enhanced dopamine responses to the protein-containing grain pellets, relative to sucrose pellets, both elicited by reward delivery and reward collection. This grain-sucrose discrimination was not evident in either NR diet phase, indicating that it emerged specifically during the protein-restricted state.

**Figure 4.**
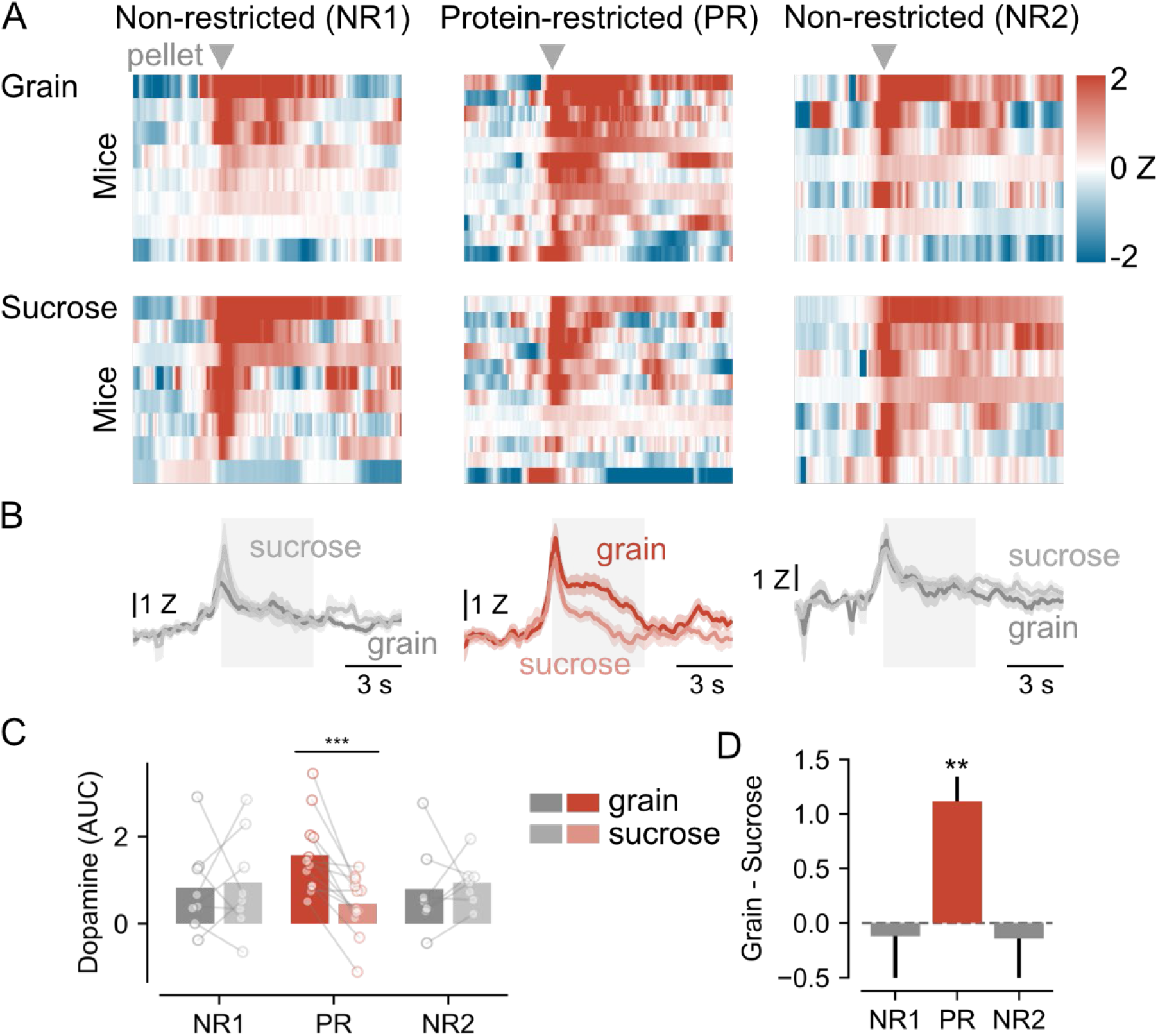
Elevated dopamine responses to grain than to sucrose when mice were protein-restricted in FR1 sessions. **(A)** Heat maps where rows are responses in individual mice to delivery of grain pellets (upper panels) and sucrose pellets (lower panels) across different phases of the experiment (non-restricted 1 → protein-restricted → non-restricted 2, left to right). Grey triangles show time of pellet delivery. (B) Line plots showing dopamine responses with thick line as mean and shaded coloured area as SEM. Shaded grey rectangle shows period averaged for area under the curve (AUC) calculation in C. (C) Dopamine AUC for grain and sucrose pellets across different experimental phases. Bars are mean and circles are individual mice. ***, p < .001 grain vs. sucrose. (D) Rectified ratio of AUC signal between grain and sucrose pellets showing that grain pellets are associated with more dopamine release than sucrose only during protein restriction phase. **, p < .01 vs. 0.5.

### 3.5 Progressive ratio (effort-based): event-specific and attenuated dopamine modulation by dietary phase

During progressive ratio, NAc dopamine responses were analysed at pellet delivery, which in these sessions marked completion of the response requirement (Fig. 5A-B). Grain and sucrose responses were compared within each dietary phase using the 0-5 s post-event AUC window (Fig. 5C). In contrast to findings under fixed ratio schedule, there was no significant effect of pellet type in any dietary phase. Grain and sucrose responses did not differ in NR1 (F_1,9_ = 0.14, p = .715), PR (F_1,9_ = 1.85, p = .206), or NR2 (F_1,5_ = 1.82, p = .235). Further analysis revealed that there was no difference between dopamine evoked by grain and sucrose pellets in any of the diet phases (Fig. 5D); Bonferroni-corrected one-sample t-test vs. zero; NR1: t_9_ = 0.66, p = 1.0; PR: t_9_ = 1.45, p = .541; NR2: t_5_ = 0.90, p = 1.0). A similar pattern was observed for signals aligned to pellet retrieval (Fig. S5). As with FR1 schedule, latencies to retrieve pellets were typically short (Fig. S6; 97.69% ProgRatio latencies < 5 s). Retrieval latencies did not differ between grain and sucrose pellets during the NR1 phase (unpaired t-test: t_20_ = 1.19, p = .248) or within the PR phase (t_19_ = 0.93, p = .362) but were slightly shorter for sucrose pellets than grain pellets during the NR2 phase (t_13_ = 2.24, p = .043). Thus, under progressive ratio conditions, dopamine responses did not show significant pellet-specific modulation at either pellet onset or pellet offset in any dietary phase.

**Figure 5.**
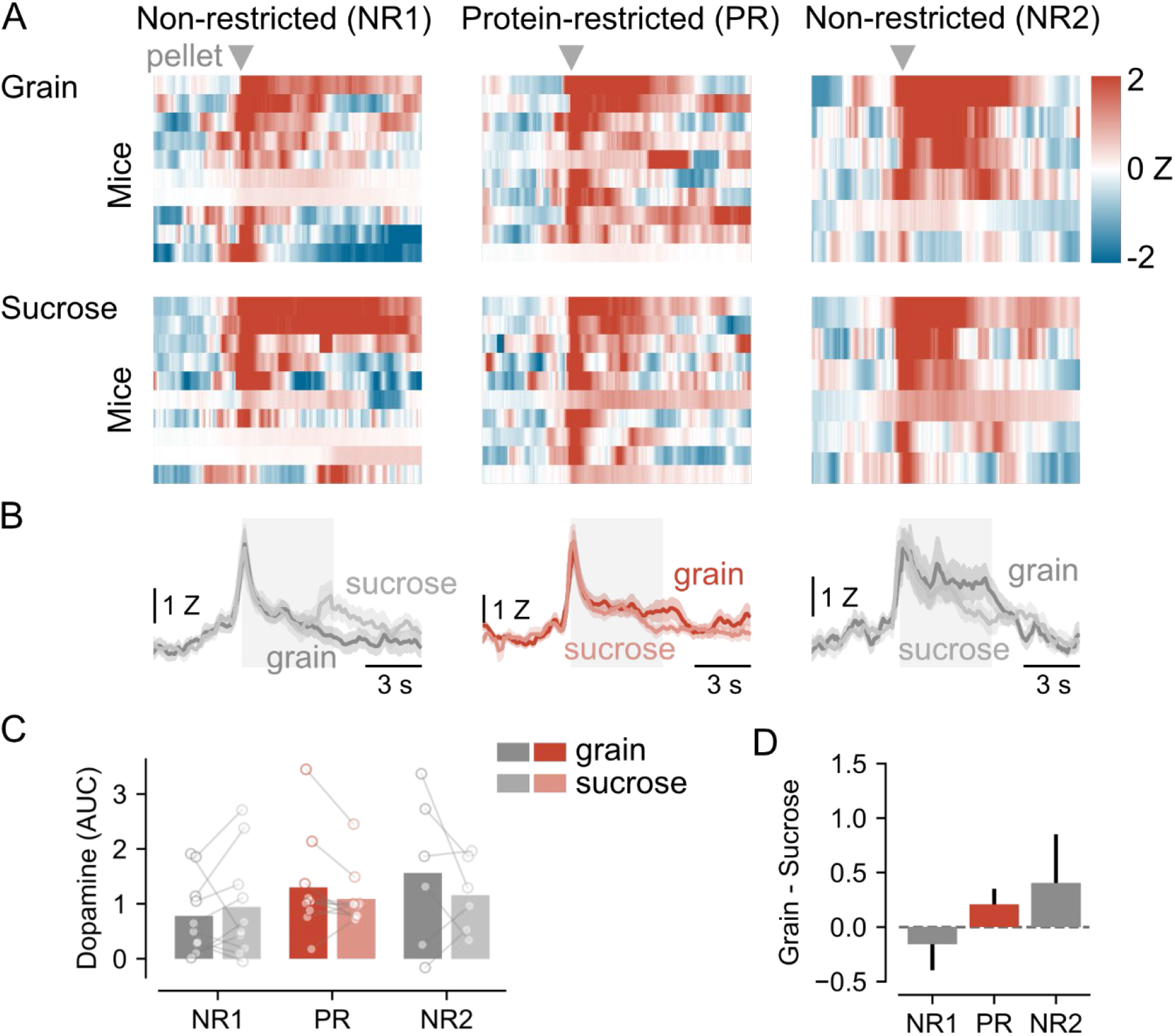
No difference in dopamine signals between grain and sucrose during progressive ratio sessions. **(A)** Heat maps where rows are responses in individual mice to delivery of grain pellets (upper panels) and sucrose pellets (lower panels) across different phases of the experiment (non-restricted 1 → protein-restricted → non-restricted 2, left to right). (B) Line plots showing dopamine responses with thick line as mean and shaded coloured area as SEM. Shaded grey rectangle shows period averaged for area under the curve (AUC) calculation in C. (C) Dopamine AUC for grain and sucrose pellets across different experimental phases. Bars are mean and circles are individual mice. (D) Rectified ratio of AUC signal between grain and sucrose pellets showing that there is no difference between pellet type in any of the diet phases.

These results indicate that, unlike the fixed ratio condition, progressive ratio sessions did not reveal significant dietary-phase-dependent discrimination between the protein-containing grain pellet and sucrose at either reward delivery or consumption. This suggests that the robust pellet-specific enhancement observed during protein restriction under low-effort conditions was attenuated or absent when rewards were earned under higher response costs. This pattern suggests that increasing effort requirement may attenuate the pellet-specific differences in dopamine signalling, such that differences evident under low-cost intake are no longer robustly expressed at reward delivery or retrieval.

## 4. Discussion

Dietary protein restriction altered NAc dopamine signalling during feeding in a manner that depended on both nutrient context and task demands. Across the three dietary phases, operant performance and dopamine did not change in parallel. Behaviourally, under FR1 and ProgRatio, protein restriction increased responding for both grain and sucrose pellets. In contrast, the clearest neural effect emerged under FR1, where NAc dopamine responses became selectively larger for grain than for sucrose during the protein-restricted phase, both at action initiation/pellet delivery and at pellet retrieval. These effects on dopamine signalling were not seen, however, during ProgRatio responding. Thus, protein restriction did not simply produce a global amplification of reward-related signalling but selectively altered NAc dopamine encoding in a state- and task-dependent manner.

Metabolic findings help define the physiological context in which these neural effects emerged. Protein restriction did not significantly increase non-FED food intake across phases, but it markedly suppressed bodyweight gain, with partial recovery after returning to the non-restricted diet. This pattern is consistent with a broader literature showing that low-protein diets can produce substantial metabolic adaptation even when changes in total energy intake are modest or variable. In particular, dietary protein restriction robustly elevates circulating FGF21, which functions as an endocrine signal of protein insufficiency and contributes to the metabolic and behavioural consequences of low-protein feeding (Laeger et al., 2014, 2016). These consequences include altered energy expenditure and reduced body weight or adiposity, indicating that the consequences of protein dilution extend beyond food intake (Hill et al., 2022; Pezeshki & Chelikani, 2021). Within that framework, the present intake and body-weight data support the conclusion that diet manipulation induced a meaningful protein-restricted physiological state against which the dopamine findings should be interpreted.

The contrast between FR1 and progressive ratio is informative. Under FR1, the low response requirement may allow nutrient-related differences in reward prediction, action initiation, and reward evaluation to be expressed more clearly in dopamine signals aligned to discrete feeding events. Under progressive ratio, by contrast, increasing effort requirements, produce longer and more variable action sequences, and lower event availability may reduce the detectability of nutrient-specific differences at pellet delivery and retrieval. This interpretation is broadly consistent with evidence that NAc circuitry contributes to effort related responding for food and to balancing feeding against competing behavioural demands (Walle et al., 2024). In that sense, the absence of significant grain-sucrose differences under progressive ratio does not necessarily imply that protein-specific motivational processes were absent, but rather that they were not robustly resolved in the present event-locked analysis.

An important feature of this study is that dopamine was recorded while mice performed in their own home cages using solid 20 mg pellets rather than flavoured nutrient solutions in a conventional operant chamber. Much of the protein-appetite literature has relied on protein (casein) and carbohydrate solutions or suspensions, which are powerful for isolating nutrient choice and post-ingestive learning, but differ substantially from pellet seeking, retrieval, and consumption in a more naturalistic environment (Chiacchierini et al., 2021; Khan et al., 2025). Our findings therefore extend existing work by showing that protein state can modulate mesolimbic dopamine signalling during interaction with solid foods in a home-cage context. At the same time, this design introduces behavioural complexity that is largely absent from standard operant boxes; mice could move freely, drink, explore, and engage with other features of the cage between feeding events. The recorded dopamine signal should therefore be understood as feeding-related activity embedded within a richer behavioural stream, which likely increases ecological validity but may also add state and context dependent variance around event-aligned responses (Taghipourbibalan & McCutcheon, 2026b).

Several limitations should be considered. First, recordings were obtained during relatively brief 1h sessions, which capture discrete feeding events well but may underestimate slower or cumulative effects of protein state on behaviour and dopamine signalling across longer timescales. Second, our analyses focused on selected event alignments and a defined post-event window, and therefore may not capture more extended task-state dynamics or interactions between feeding and other ongoing home-cage behaviours. Third, event availability varied across animals and session types, particularly during progressive ratio testing where some mice showed relatively few pellet retrieval events. Consequently, statistical power to detect subtle nutrient-specific differences in dopamine signalling may have been reduced in some analyses. Future work could therefore combine longer sessions, more temporally layered analyses, and complementary task designs to determine how protein state shapes dopamine signalling across both discrete reward events and broader motivational states. It will also be important to test more directly how endocrine signals such as FGF21 and upstream hypothalamic circuits contribute to the protein state-dependent modulation of NAc dopamine observed here.

## Author contributions

Conceptualization: HT, JEM; Methodology: HT; Formal analysis: HT, JEM; Investigation: HT, SWH, KLV; Writing – Original draft: HT; Writing – Review and editing: JEM, KLV; Supervision: JEM, KLV; Funding acquisition: JEM.

## Acknowledgements

The authors would like to acknowledge the staff at the Department of Comparative Medicine (AKM) and the associated animal facility of the Faculty of Health Sciences at UiT and the staff working in the workshops of the Faculty of Health Sciences at UiT.

