## Supplementary Material for "Protein restriction amplifies nucleus accumbens dopamine responses to protein-containing food during operant feeding in male mice"

The figures, tables, and supplementary sections in this document are arranged in the order of their first appearance in the main manuscript.

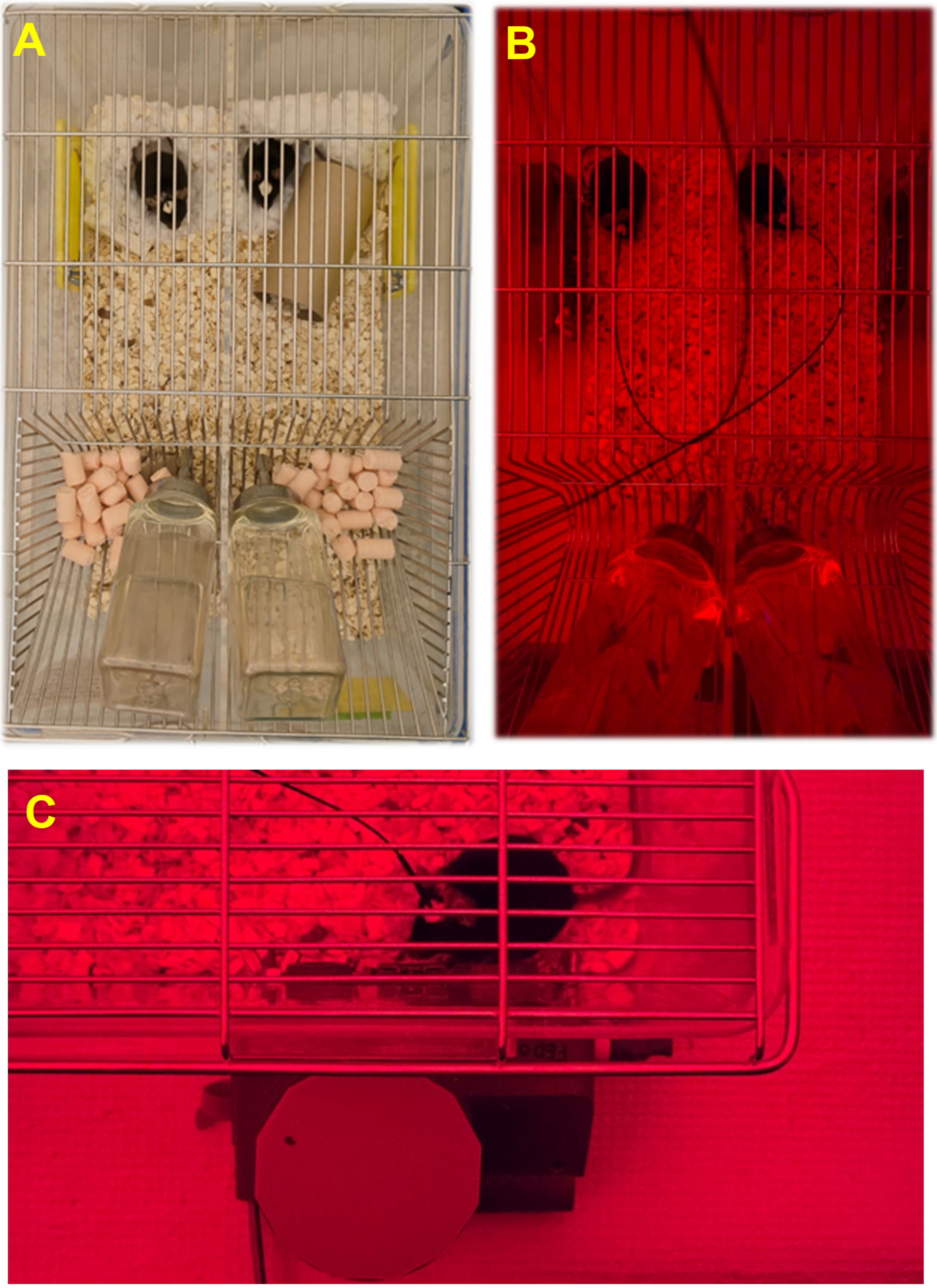


**Fig. S1. Modified FED3-compatible home cage used throughout the study.** Mice were housed two per cage in modified FED3-compatible cages separated by a perforated divider that permitted visual, olfactory, and limited tactile communication while preventing direct physical interaction. The cages contained two lateral ports designed to accommodate FED3 devices. **(A)** When FED3 devices were not in use (i.e. outside of FED3 training and fiber photometry recording sessions), the ports were sealed with removable 3D-printed covers. **(B, C)** During behavioral recording sessions, FED3 devices were mounted to the lateral ports, allowing mice to perform operant responding while remaining in their home cages.

### Table S1. Composition of diets and operant-session pellets

#### (A) Composition of the diets used for diet manipulation in the home cage

| Ingredient | 20% Casein | | | 5% Casein | | |
|---|---|---|---|---|---|---|
| | Gram | | | Gram | | |
| Casein | 200 | | | 50 | | |
| L-Cystine | 3 | | | 0.75 | | |
| Corn Starch | 375.7 | | | 485 | | |
| Maltodextrin 10 | 125 | | | 150 | | |
| Sucrose | 107.0777 | | | 107.0777 | | |
| Cellulose | 50 | | | 50 | | |
| Soybean Oil | 25 | | | 25 | | |
| Lard | 75 | | | 75 | | |
| Mineral Mix S10022G | 0 | | | 0 | | |
| Mineral Mix S10022C | 3.5 | | | 3.5 | | |
| Calcium Carbonate | 12.495 | | | 8.7 | | |
| Calcium Phosphate, Dibasic | 0 | | | 5.3 | | |
| Potassium Citrate, 1 $H_2O$ | 2.4773 | | | 2.4773 | | |
| Potassium Phosphate, Monobasic | 6.86 | | | 6.86 | | |
| Sodium Chloride | 2.59 | | | 2.59 | | |
| Vitamin Mix V10037 | 10 | | | 10 | | |
| Choline Bitartrate | 2.5 | | | 2.5 | | |
| FD&C Yellow Dye #5 | 0.05 | | | 0 | | |
| FD&C Red Dye #40 | 0 | | | 0.05 | | |
| FD&C Blue Dye #1 | 0 | | | 0 | | |
| **Total** | **1001.250** | | | **984.805** | | |
| **Nutrient** | **gm** | **kcal** | **gm% / kcal%** | **gm** | **kcal** | **gm% / kcal%** |
| Protein | 179 | 716 | 18 / 18 | 45 | 179 | 5 / 4 |
| Carbohydrate | 618 | 2471 | 62 / 60 | 752 | 3008 | 76 / 74 |
| Fat | 100 | 900 | 10 / 22 | 100 | 900 | 10 / 22 |
| Fiber | 50 | 0 | – | 50 | 0 | – |
| **Total kcal** | **4087** | | | **4087** | | |
| **Measure** | **20% Casein** | | | **5% Casein** | | |
| Calcium (g) | 5.06 | | | 5.06 | | |
| Phosphorus (g) | 3.16 | | | 3.17 | | |
| Potassium (g) | 3.60 | | | 3.60 | | |
| Folate (mg) | 2.0 | | | 2.0 | | |

**(B) Composition of the 20 mg grain pellets used during operant FED3 sessions**

#### Proximate Profile

| | | |
|---|---|---|
| Protein | % | 21.3 |
| Fat | % | 3.8 |
| Fiber | % | 4.0 |
| Ash | % | 8.1 |
| Moisture | % | $< 10$ |
| Carbohydrate | % | 54.0 |

#### Caloric Profile

| | | |
|---|---|---|
| Protein | kcal/gm | 0.85 |
| Fat | kcal/gm | 0.34 |
| Carbohydrate | kcal/gm | 2.16 |
| **Total** | kcal/gm | **3.35** |

#### Amino Acids

| | | |
|---|---|---|
| Alanine | gm/kg | 11.0 |
| Arginine | gm/kg | 10.8 |
| Aspartic Acid | gm/kg | 17.4 |
| Cystine | gm/kg | 2.8 |
| Glutamic Acid | gm/kg | 37.9 |
| Glycine | gm/kg | 8.5 |
| Histidine | gm/kg | 5.7 |
| Isoleucine | gm/kg | 10.2 |
| Leucine | gm/kg | 21.0 |
| Lysine | gm/kg | 11.0 |
| Methionine | gm/kg | 7.5 |
| Phenylalanine | gm/kg | 10.6 |
| Proline | gm/kg | 16.7 |
| Serine | gm/kg | 11.3 |
| Threonine | gm/kg | 8.4 |
| Tryptophan | gm/kg | 2.2 |
| Tyrosine | gm/kg | 8.1 |
| Valine | gm/kg | 11.5 |

#### Carbohydrates

| | | |
|---|---|---|
| Monosaccharides | gm/kg | 105 |
| Disaccharides | gm/kg | 140 |
| Polysaccharides | gm/kg | 278 |

#### Fatty Acids

| | | |
|---|---|---|
| C18:2 Linoleic | gm/kg | 14.7 |
| C18:3 Linolenic | gm/kg | 1.4 |
| Total Saturated | gm/kg | 8.2 |
| Total Monounsaturated | gm/kg | 10.6 |
| Total Polyunsaturated | gm/kg | 16.3 |

#### Minerals

| | | |
|---|---|---|
| Calcium | gm/kg | 11.2 |
| Chloride | gm/kg | 6.6 |
| Copper | mg/kg | 18.9 |
| Iodine | mg/kg | 6.2 |
| Iron | mg/kg | 430 |
| Magnesium | gm/kg | 1.9 |
| Manganese | mg/kg | 53.8 |
| Phosphorus | gm/kg | 10.7 |
| Potassium | gm/kg | 11.3 |
| Selenium | mg/kg | 0.23 |
| Sodium | mg/kg | 3413 |
| Zinc | mg/kg | 75.9 |

#### Vitamins

| | | |
|---|---|---|
| Ascorbic Acid | mg/kg | 1491 |
| Biotin | mg/kg | 0.41 |
| Choline | mg/kg | 2414 |
| Folate | mg/kg | 11.4 |
| Niacin | mg/kg | 96.6 |
| Pantothenic Acid | mg/kg | 54.9 |
| Pyridoxine | mg/kg | 32.3 |
| Riboflavin | mg/kg | 31.9 |
| Thiamin | mg/kg | 31.4 |
| Vitamin A | IU/kg | 19788 |
| Vitamin $B_{12}$ | mcg/kg | 50 |
| Vitamin $D_3$ | IU/kg | 5757 |
| Vitamin E | IU/kg | 136 |
| Vitamin $K_3$ (Menadione) | mg/kg | 48.5 |

#### Ingredients

Ground Corn, Banana Flakes, Dehulled Soybean Meal, Corn Gluten Meal, Ground Wheat, Corn Gluten Feed, Fish Meal, Dehydrated Alfalfa Meal, Casein, Dried Whey, Sucrose, Fructose, Dextrose, Dried Beet Pulp, Soybean Oil, Porcine Animal Fat (preserved with BHA), Dried Brewers Yeast, Mineral Mix, Vitamin Mix, Magnesium Stearate, DL-Methionine, Choline Chloride, Sodium Propionate, Ascorbic Acid, Kaolin.

**(C) Composition of the 20 mg sucrose pellets used during operant FED3 sessions**

| Proximate Profile | | |
|---|---|---|
| Protein | % | 0.0 |
| Fat | % | 0.5 |
| Fiber | % | 0.0 |
| Ash | % | 1.5 |
| Moisture | % | $< 5$ |
| Carbohydrate | % | 94.8 |
| **Caloric Profile** | | |
| Protein | kcal/gm | 0.0 |
| Fat | kcal/gm | 0.05 |
| Carbohydrate | kcal/gm | 3.79 |
| **Total** | kcal/gm | **3.84** |
| **Carbohydrates** | | |
| Monosaccharides | gm/kg | 348 |
| Disaccharides | gm/kg | 600 |
| Polysaccharides | gm/kg | 0 |

**Ingredients**

Sucrose, Dextrose, Magnesium Stearate, Calcium Silicate, Mineral Oil.

#### S2.5.2. Fibre photometry signal processing and statistical analysis

##### Photometry acquisition and preprocessing

Fibre photometry recordings were acquired using a dual-wavelength configuration consisting of an activity-dependent blue excitation channel and a simultaneously recorded UV isosbestic control channel collected through the same optical fibre. Raw photometry data were imported from Tucker-Davis Technologies (TDT) tank files and processed in Python using custom scripts together with the `tdt` and `trompy` libraries.

For each recording session, the blue and UV streams were extracted from the TDT block and processed using the Trompy signal-processing pipeline. The UV signal was used to correct the blue signal for shared non-neural variance, including motion-related artefacts and slow drift, and the corrected signal was z-scored within session. All subsequent analyses were therefore performed on motion-corrected, session-normalised photometry traces.

##### Behavioural event definition and peri-event extraction

Behavioural events were identified from TTL pulses recorded concurrently with the behavioural session. The event-extraction pipeline was configured to detect left-poke onset, right-poke onset, pellet onset, and pellet offset TTLs. The analyses reported here focused on event families relevant to the final behavioural design: FR1 left-poke(active rewarding poke) onset, FR1 pellet retrieval (pellet offset), progressive-ratio pellet delivery (pellet onset), and progressive-ratio pellet retrieval (pellet offset).

For each valid TTL event, a peri-event photometry segment was extracted from $-5\,\mathrm{s}$ to $+10\,\mathrm{s}$ relative to the event timestamp. Segments were sampled at the native acquisition frequency of the recording, producing uniformly sampled peri-event traces. If a required TTL channel was absent or contained no valid events, that recording was excluded from the corresponding event-specific analysis.

##### Event-level storage in flexible pickle files

To retain flexibility for downstream analysis, extracted peri-event traces were first stored at the event level. For each unique combination of session number, diet, pellet type, behavioural schedule, event type, and alignment point, all peri-event segments were saved in a condition-specific pickle file. Each file contained the full matrix of peri-event traces together with the corresponding event times, mouse IDs, session numbers, event indices within session, and the shared peri-event time vector.

In addition, each pickle file included a descriptive event-level summary trace with pointwise confidence intervals generated by bootstrap resampling across events. These summaries were retained for convenience and visual inspection only and were not used for formal statistical inference.

##### Rationale for restricting analyses to the first $K$ events per mouse

Mice did not necessarily obtain the same number of pellets or make the same number of responses across sessions, pellet types, or behavioural schedules. Moreover, later events within a session may reflect a different behavioural or neural state from earlier events. To standardise event sampling across mice and conditions, inferential analyses were therefore restricted to the first $K$ events per mouse within each condition. Event order was determined using the stored within-session event index when available, and otherwise by event timestamp. Mice with fewer than $K$ valid events in a given condition were excluded from that specific analysis.

##### Behavioural event-availability analysis used to define $K$

The value of $K$ was determined from a separate behavioural event-availability analysis performed prior to the final photometry statistics. Using the behavioural summary data, the number of relevant events per mouse was quantified for each combination of phase, pellet type, behavioural mode, and event family. For a range of candidate thresholds, survival-style plots were generated showing how many mice retained at least $K$ events under each condition.

The purpose of this procedure was not merely to maximise sample size, but to identify a threshold that preserved a reasonable number of mice while ensuring that the compared photometry signals were derived from similarly early events in each session. Based on these distributions, the final thresholds were set as follows:

- FR1 left-poke onset/offset: $K = 6$
- Prog-ratio pellet onset/offset: $K = 4$

These thresholds provided a compromise between retaining sufficient mice in each condition and restricting analysis to an early and reasonably matched portion of each session.

##### Mouse-level averaging and baseline correction

All inferential analyses were conducted at the level of the mouse rather than the individual event. After first-$K$ event selection, the retained peri-event segments were averaged within each mouse to generate a single mean peri-event trace for each mouse and condition. Each mouse-level trace was then baseline-corrected by subtracting the mean signal during the $-2$ to $0\,\mathrm{s}$ pre-event interval.

##### Quantification of event-evoked responses

Event-evoked responses were quantified as the area under the baseline-corrected peri-event trace during the $0$ to $5\,\mathrm{s}$ post-event window. AUC was calculated numerically using trapezoidal integration (`numpy.trapz`).

##### Experimental conditions and comparison structure

The final reported analyses focused on Grain and Sucrose sessions across the three diet phases: NR1, PR, and NR2. For each phase, the relevant behavioural sessions were FR1 Grain, progressive-ratio Grain, FR1 Sucrose, and progressive-ratio Sucrose, with event-specific analyses performed separately for FR1 left-poke onset, FR1 pellet offset, progressive-ratio pellet onset, and progressive-ratio pellet offset. The primary comparisons were performed within each phase, so dopamine signals of responses to Grain vs Sucrose were compared with each other in each phase.

##### Statistical analysis

All statistical analyses were performed in Python using mouse-level AUC values derived from the baseline-corrected traces. Mouse identity was treated as the repeated-measures unit throughout.

For each event family, Grain and Sucrose responses were compared separately within each phase. Specifically, analyses were carried out for:

- FR1 left-poke onset
- FR1 pellet offset
- Progressive-ratio pellet onset
- Progressive-ratio pellet offset

Within each phase, a one-factor repeated-measures ANOVA was performed with pellet type (Grain, Sucrose) as the within-subject factor. Paired $t$-tests were then used as planned within-phase contrasts between Grain and Sucrose. Holm correction was applied across the three phase-specific post hoc pellet comparisons within each event family. In the plotting workflow, significance was annotated only when the omnibus repeated-measures ANOVA was significant and the Holm-corrected paired comparison also reached significance.

##### Completeness and exclusion criteria

Only mice with complete data for both pellet conditions within a given phase were included in the corresponding repeated-measures analysis. Consequently, the set of mice included could vary across event families and phases depending on event availability and the first-$K$ filtering step. Any predefined mouse exclusions listed in the exclusion file were applied before event selection and statistical analysis.

##### Visualisation of peri-event traces

For plot presentation, traces were also summarised at the mouse level after first-$K$ selection and baseline correction. Group mean peri-event traces were plotted together with $95\%$ confidence intervals generated by bootstrap resampling across mice. Thus, visualisation preserved the mouse as the biological unit of replication.

Within each phase, figures displayed baseline-corrected Grain and Sucrose traces overlaid for the relevant event type, together with paired mouse-level AUC scatter plots. Significance annotations shown on the figures were based on the Holm-corrected mouse-level analyses described above.

##### Relationship between event-level extraction, mouse-level summarisation, and inference

The overall analysis pipeline was intentionally separated into distinct stages. First, all available peri-event signals were extracted and stored at the event level in flexible pickle files. Second, event inclusion rules were applied to select the first $K$ events per mouse, after which traces were averaged within mouse and baseline-corrected. Third, formal statistical analyses were performed on mouse-level AUC values derived from these baseline-corrected mean traces.

This structure ensured that individual events were not treated as independent biological replicates in inferential statistics, while still preserving complete event-level data for transparency, quality control, and flexible visualisation.

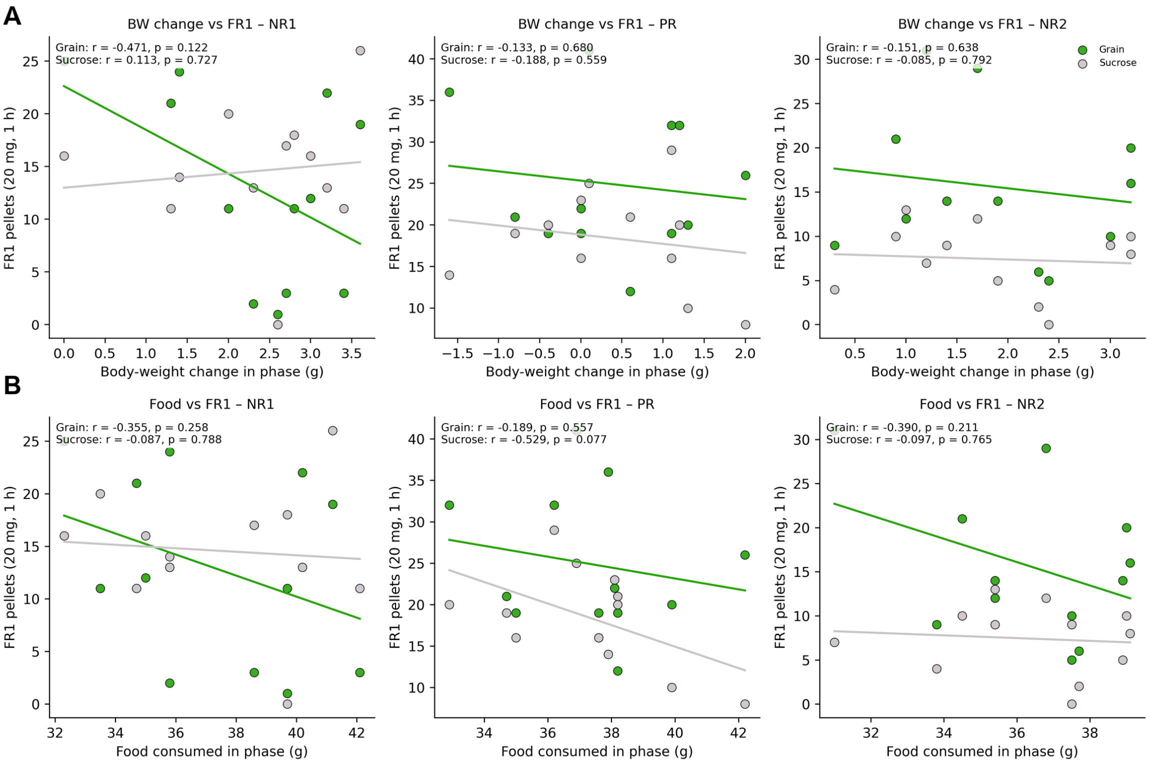


**Fig. S2. Inter-individual variability in operant responding is not explained by energy balance.** Pearson correlation analyses were performed separately for each dietary phase and pellet type. **(A)** Correlations between FR1 pellet count and bodyweight change measured across the corresponding dietary phase for grain and sucrose pellets. **(B)** Correlations between FR1 pellet count and home-cage food intake measured across the corresponding dietary phase for grain and sucrose pellets. No significant correlations were detected between FR1 pellet count and either bodyweight change or home-cage food intake in any dietary phase for either grain or sucrose pellets.

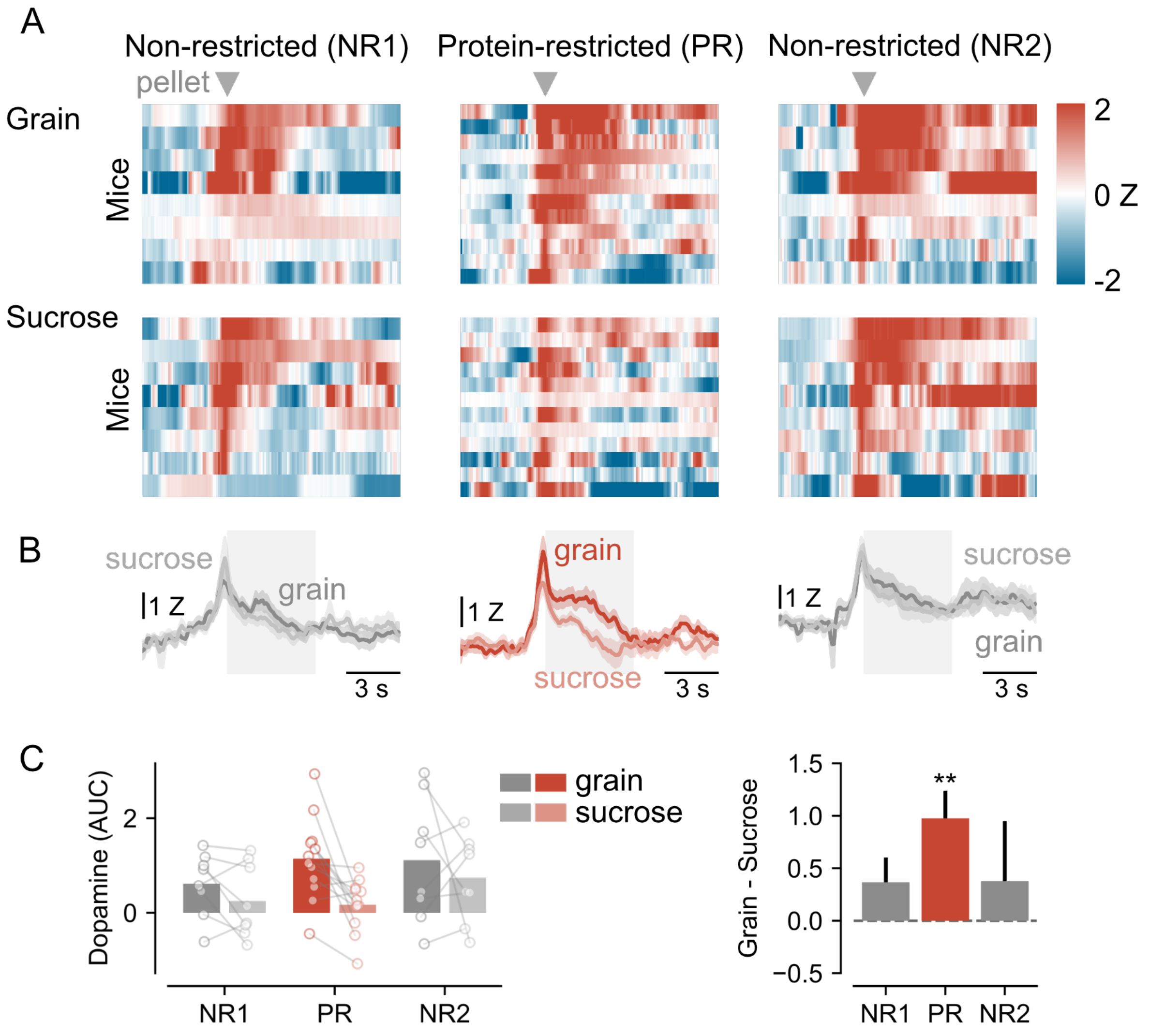


**Fig. S3. Pellet retrieval-aligned nucleus accumbens dopamine responses under FR1. (A)** Heat maps of z-scored dopamine signals aligned to pellet retrieval (time 0) for individual mice during the first non-restricted (NR1), protein-restricted (PR), and second non-restricted (NR2) dietary phases. Grain and sucrose sessions are shown separately. Each row represents one mouse. **(B)** Mean ± SEM dopamine traces aligned to pellet retrieval for grain and sucrose pellets in each dietary phase. The shaded region (0–5 s) indicates the time window used for area-under-the-curve (AUC) quantification. **(C)** Left, 0–5 s post-retrieval dopamine AUC for grain and sucrose pellets across dietary phases. During PR, dopamine responses at pellet retrieval were significantly greater for grain than sucrose ($F(1,11) = 11.10$, $p = 0.0067$; Holm-corrected post hoc $p = 0.0201$), whereas no significant pellet-type difference was found in NR1 ($F(1,7) = 2.55$, $p = 0.154$) or NR2 ($F(1,6) = 0.003$, $p = 0.959$). Right, difference in dopamine AUC (grain – sucrose) across dietary phases. Bonferroni-corrected one-sample t-test vs. zero; NR1: $t7 = 1.55$, $p = .493$; PR: $t11 = 3.72$, $p = .010$; NR2: $t7 = 0.66$, $p = 1.0$).

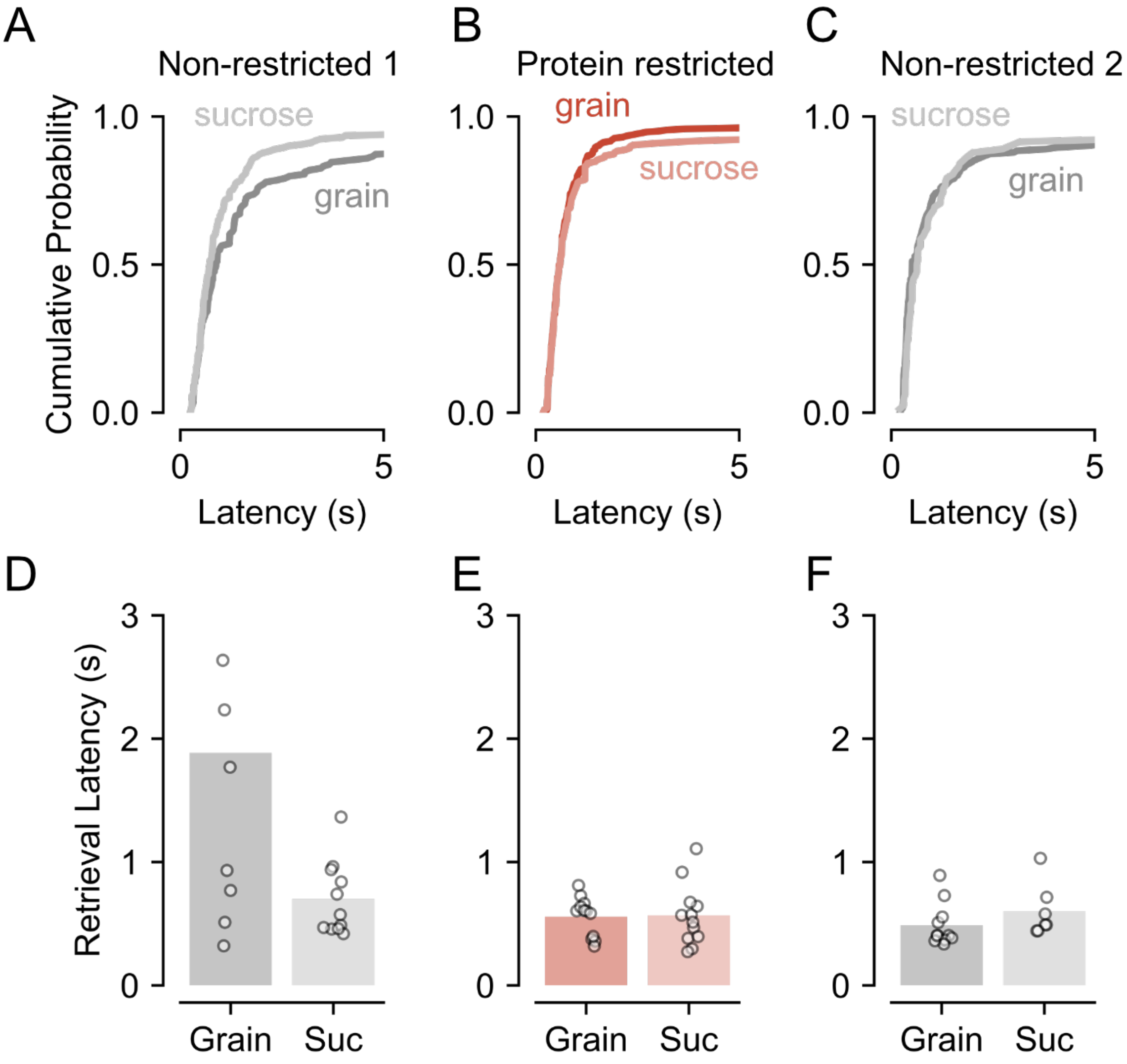


**Fig. S4. Pellet retrieval latencies under FR1 across dietary phases. (A–C)** Cumulative distributions of pellet retrieval latencies following pellet delivery for grain and sucrose pellets during the first non-restricted (NR1), protein-restricted (PR), and second non-restricted (NR2) dietary phases, respectively. Across all phases, the majority of pellets were retrieved within the first few seconds after delivery. **(D–F)** Median pellet retrieval latency for grain and sucrose pellets during NR1, PR, and NR2, respectively. Retrieval latencies did not differ significantly between grain and sucrose pellets within any dietary phase, indicating that differences in dopamine responses were unlikely to be explained by differences in reward retrieval timing.

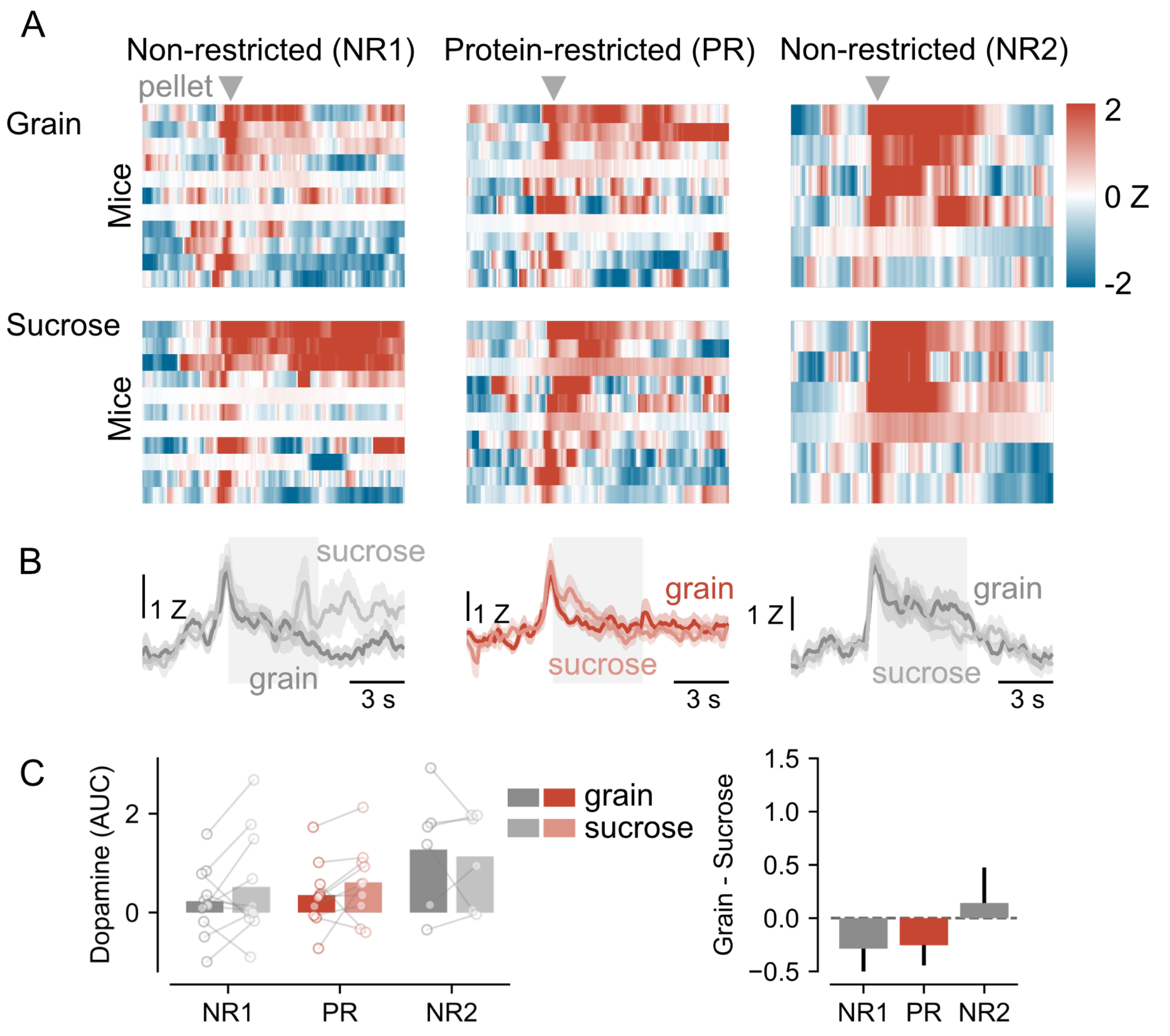


**Fig. S5. Pellet retrieval-aligned nucleus accumbens dopamine responses under progressive ratio. (A)** Heat maps of z-scored dopamine signals aligned to pellet retrieval (time 0) for individual mice during the first non-restricted (NR1), protein-restricted (PR), and second non-restricted (NR2) dietary phases. Grain and sucrose sessions are shown separately. Each row represents one mouse. **(B)**Mean ± SEM dopamine traces aligned to pellet retrieval for grain and sucrose pellets in each dietary phase. The shaded region (0–5 s) indicates the time window used for area-under-the-curve (AUC) quantification. **(C)** Left, 0–5 s post-retrieval dopamine AUC for grain and sucrose pellets across dietary phases. Consistent with pellet delivery-aligned responses (Fig. 5), retrieval-aligned dopamine responses did not differ significantly between grain and sucrose pellets in NR1 ($F(1,10) = 0.005$, $p = 0.947$), PR ($F(1,9) = 0.001$, $p = 0.979$), or NR2 ($F(1,5) = 0.94$, $p = 0.376$). Right, difference in dopamine AUC (grain – sucrose) across dietary phases. Bonferroni-corrected one-sample t-test vs. zero; NR1: $t10 = 1.26$, $p = .708$; PR: $t9 = 1.33$, $p = .653$; NR2: $t5 = 0.43$, $p = 1.0$).

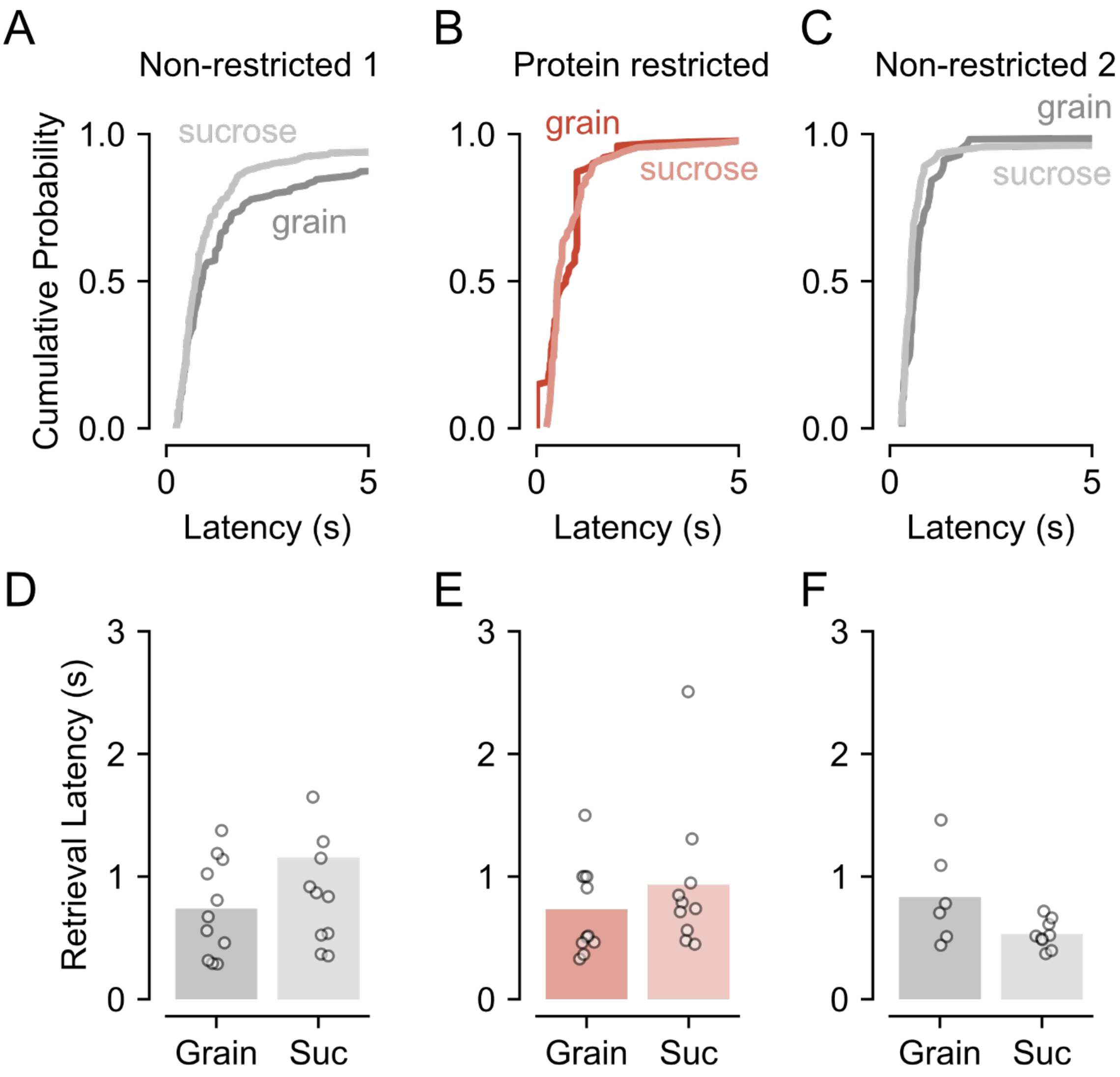


**Fig. S6. Pellet retrieval latencies under progressive ratio across dietary phases. (A–C)** Cumulative distributions of pellet retrieval latencies following pellet delivery for grain and sucrose pellets during the first non-restricted (NR1), protein-restricted (PR), and second non-restricted (NR2) dietary phases, respectively. Across all phases, the majority of pellets were retrieved within the first few seconds after delivery. **(D–F)**Median pellet retrieval latency for grain and sucrose pellets during NR1, PR, and NR2, respectively. Retrieval latencies did not differ significantly between grain and sucrose pellets within any dietary phase, indicating that differences in dopamine responses were unlikely to be explained by differences in reward retrieval timing.
